# The b^0+^ Amino Acid Transporter Defines a Selenium-utilization Enterocyte Program in the Human Intestine

**DOI:** 10.64898/2026.05.27.728150

**Authors:** Xiaobai He, Kundu Zhong, Wangzhen Yang, Jiahao Cao, Xiaopan Chen, Linjie Chen

## Abstract

Selenium is an essential trace element incorporated into selenoproteins as selenocysteine, yet the intestinal cellular programs associated with selenium utilization remain poorly defined. Here, we performed a large-scale single-cell transcriptomic analysis of 169,068 human enterocytes from 105 donors to systematically profile nine candidate selenium transporter systems and their relationship to selenoprotein expression. Among all transporters examined, the b^0+^ amino acid transporter system (SLC3A1/SLC7A9) showed the strongest and most maturation-independent association with selenoproteins, including SELENOP (Spearman *r* = 0.489) and GPX4 (*r* = 0.392), with enrichment across 21 of 24 detected selenoproteins in b^0+^-complete versus transporter-negative enterocytes. These associations were robust across individual donors and confirmed by pseudobulk validation, and were only partially explained by enterocyte maturation state, intestinal segment, and sequencing depth after covariate adjustment, indicating both differentiation-dependent and differentiation-independent components. Furthermore, enterocytes carrying the b^0+^ complex expressed a non-canonical LRP repertoire (LRP1, LRP5 and LRP6) rather than the canonical SELENOP receptors LRP2 and LRP8; this co-expression was maturation-dependent, nominating these LRPs as candidate intestinal SELENOP-handling receptors. Together, these single-cell data identify b^0+^ transporter expression as a marker of a selenoprotein-enriched enterocyte state in the human intestine.

## Introduction

Selenium is an essential micronutrient whose biological functions are mediated primarily through its cotranslational incorporation into selenoproteins as selenocysteine^1^. The human selenoproteome comprises 25 members with critical roles in antioxidant defense, redox regulation, and cellular homeostasis, including the ferroptosis-suppressive phospholipid hydroperoxide glutathione peroxidase GPX4^1–3^ and the systemic selenium transport protein SELENOP^4,5^. Intestinal enterocytes constitute the primary site of dietary selenium absorption and thus control systemic selenoprotein biosynthetic capacity, yet the transporter systems that define selenium-utilization programs in human intestinal epithelium remain incompletely characterized^6^.

Dietary selenium exists in several chemical forms, principally organic selenomethionine and selenocysteine and inorganic selenite and selenate, each absorbed by largely distinct carrier systems in the intestinal epithelium. Free selenocysteine liberated upon proteolysis of dietary selenoproteins is chemically unstable and expected to exist predominantly as its diselenide dimer selenocystine (the Se analog of cystine) in the oxidizing milieu of the gut lumen^7–9^. The b^0+^ heterodimeric transporter (SLC3A1/SLC7A9) and the cystine-glutamate antiporter xCT (SLC3A2/SLC7A11) have both been proposed as candidate selenocystine transporters^10–12^, yet their relative importance in normal human absorptive enterocytes has not been directly compared. Direct functional evidence for b^0+^-mediated selenocystine transport has recently been obtained in a companion study from our group, demonstrating saturable, SLC3A1/SLC7A9-dependent uptake with substrate selectivity consistent with canonical b^0+^ transport^13^. The cellular context and physiological relevance of this transport activity in primary human intestinal epithelium, however, remain undefined, a question we address in the present study. More broadly, no study has systematically co-profiled candidate selenium transporter systems alongside selenoprotein expression programs across human intestinal cell types at single-cell resolution, a gap that prior amino acid transporter atlases, focused on general nutrient absorption, did not address^14^.

Recent aggregation of large-scale human intestinal single-cell transcriptomic datasets in resources such as the CellxGene Census now enables systematic interrogation of transporter expression across diverse cell populations, intestinal segments, and donor backgrounds at a scale not previously achievable^15,16^. Critically, concurrent measurement of selenoprotein expression in the same cells allows correlative analysis linking candidate transporter programs to potential downstream selenium utilization states, an approach that has not been applied to normal human intestinal epithelium. However, interpreting such correlations requires explicit treatment of a major potential confound: enterocyte differentiation state co-varies with multiple metabolic gene programs, necessitating confound-aware analytical strategies to distinguish selenium-specific associations from generic absorptive maturation signals^17–20^.

Here, we performed a large-scale single-cell transcriptomic analysis of 169,068 primary human intestinal enterocytes from 105 donors, systematically comparing nine candidate selenium transporter systems representing major dietary selenium chemical forms against 24 of the 25 annotated human selenoproteins. Using partial correlation analyses controlling for enterocyte maturation state and intestinal segment identity, combined with donor-stratified validation, we identify the b^0+^ transporter system (SLC3A1/SLC7A9) as the system most robustly associated with a coordinated selenoprotein-enriched enterocyte transcriptional state, an association that persists after adjustment for differentiation-related confounders and is reproducible across individuals. These findings provide a confound-aware, systems-level transcriptomic framework for selenium utilization in the human intestinal epithelium.

## Methods

### 1. Data source, cell selection, and quality control

Single-cell RNA-sequencing data were obtained from the CellxGene Census (version 2025-11-08), a harmonized repository of publicly available human single-cell datasets, accessed programmatically with the cellxgene-census Python package. Three intestinal datasets with granular segment annotation were queried by dataset identifier (Table S4)^21,22^, restricted to Homo sapiens and primary data (is_primary_data == True). Expression matrices were retrieved in AnnData format with cell metadata including tissue, cell_type, dataset_id, assay, and donor_id. Absorptive cells were identified by case-insensitive keyword matching on the cell_type field (“enterocyte”, “colonocyte”, “colon epithelial”, “absorptive”, “gut absorptive”, “BEST4”), excluding secretory, stromal, and immune populations; this yielded 169,068 enterocytes (12.3% of 1,370,011 intestinal cells) from 105 donors across the three datasets. M cells of Peyer’s patches (cell_type containing “M cell”) were excluded a priori from all analyses, as their antigen-sampling program is not part of the absorptive selenium-uptake physiology under study. We used the Census-harmonized expression matrices as provided; quality control performed by the originating studies and by the Census pipeline was therefore inherited, and no additional per-cell filtering (e.g., on counts, detected genes, or mitochondrial fraction) was applied beyond the tissue- and cell-type selection described above. To ensure that this choice and any residual differences in capture efficiency did not drive the reported associations, sequencing depth was explicitly modeled as a covariate.

### 2. Gene sets and expression quantification

Five gene sets were retrieved and merged by cell barcode (full lists in Supplementary Methods): 11 transporter subunits spanning nine candidate selenium transporter systems; 24 of the 25 annotated human selenoproteins (SELENOF/SEP15 was absent from the Census gene index and could not be included); five selenocysteine-insertion machinery genes; five LRP-family receptors; and 10 canonical absorptive maturation markers. Ensembl IDs were mapped to HGNC symbols via the feature_name field. All expression values are raw read counts as stored in the Census; no library-size normalization was applied. This choice is appropriate because the primary statistics are rank-based (Spearman correlation, which is invariant to any monotonic per-cell scaling) or threshold-based (detection rate). The detection rate of a gene in a cell population was defined as the fraction of cells with count > 0, and mean expression as the arithmetic mean of raw counts across all cells (including zeros). For count-based summaries that are not rank-invariant—group mean expression and per-donor pseudobulk means—robustness to between-cell and between-donor depth differences was confirmed by the depth-sensitivity analyses.

### 3. Transporter classification and three-group stratification

For heterodimeric systems, transcript-level co-expression required simultaneous detection (count > 0) of both heavy- and light-chain subunits; for monomeric systems, detection of the single subunit was used. Because the SLC3A2 heavy chain is shared by xCT, LAT1, and LAT2, cells were classified independently for each system by their respective light chain. For b^0+^-specific analyses, enterocytes were partitioned into three mutually exclusive groups: b^0+^-complete (SLC3A1^+^/SLC7A9^+^, n = 21,234), SLC7A9-only (SLC3A1^−^/SLC7A9^+^, n = 11,874), and negative (SLC3A1^−^/SLC7A9^−^, n = 106,961). Cells expressing SLC3A1 alone (n = 28,999) were excluded from three-group analyses, as this configuration is not expected to form a functional transporter.

### 4. Differential expression and enrichment statistics

Three-group comparisons used the Mann-Whitney U test for all three pairwise combinations (b^0+^-complete vs. negative; b^0+^-complete vs. SLC7A9-only) and, where all three groups were assessed jointly, the Kruskal–Wallis test. Log_2_ fold change was computed as log_2_[(mean_pos + ε)/(mean_neg + ε)] with ε = 1×10^−6^; for genes with zero mean expression in both groups, log_2_FC is undefined and reported as NA. For genes with zero detection in both compared groups, the Mann-Whitney U statistic is undefined (U = n₁×n₂/2 by convention, *r* = 0.500); such entries are reported as NA for effect size and are not interpreted. A selenoprotein was scored as showing a complete monotonic gradient when detection followed b^0+^-complete > SLC7A9-only > negative with a significant b^0+^-complete vs. negative comparison. Selenoproteins with higher detection in the negative group than in b^0+^-complete (log2FC < 0) were not considered enriched in the b^0+^ program and are reported separately from the positive-gradient genes. For segment-level statistics, a minimum of 50 cells per segment was required before computing detection rates, and small- vs. large-intestine comparisons used the Mann-Whitney U test. All p-values within a given analysis were corrected across the relevant family of tests (e.g., the 9×24 transporter–selenoprotein matrix = 216 pairs) by the Benjamini-Hochberg (BH) false-discovery-rate procedure; BH-adjusted *p* < 0.05 was considered significant. Given the large cell numbers, effect sizes rather than p-values are emphasized throughout: log_2_FC as defined above; Spearman *r* (range [−1, 1]); and rank-biserial *r*, computed as *r* = U/(n_1_ × n_2_) where U is the Mann-Whitney statistic and n_1_, n_2_ are the group sizes (range [0, 1], interpretable as the probability that a randomly drawn b^0+^-complete cell exceeds a randomly drawn comparator cell in expression).

### 5. Correlation and confound-adjusted partial correlation

Spearman rank correlations and two-tailed p-values were computed for all transporter–selenoprotein pairs across the full enterocyte population (scipy.stats.spearmanr), with BH FDR correction. To separate transporter-specific signal from the enterocyte-differentiation gradient, partial Spearman correlations were computed controlling for two covariates: (i) an enterocyte maturation score, defined as the mean, across 10 canonical absorptive markers, of each marker’s within-population fractional rank (rank divided by the number of cells, giving a per-marker quantile in [0,1]); and (ii) intestinal segment, encoded as a single ordinal variable reflecting proximal-to-distal position (one-dimensional encoding was used deliberately to capture the monotonic absorptive gradient with a single degree of freedom). Partial correlations used the rank-residual method: SLC7A9 and selenoprotein expression were each rank-transformed and regressed on the covariates (intercept + covariate ranks) by ordinary least squares, and the residuals correlated by Pearson correlation; p-values were obtained from the t-distribution with n-k-2 degrees of freedom (k = number of covariates) and BH-corrected. Four covariate conditions were evaluated: unadjusted, maturation only, segment only, and fully adjusted (maturation + segment). Total *r*^2^ was decomposed into maturation-attributable, segment-attributable, shared, and residual components from pairwise differences between adjusted and unadjusted models. Within-segment validation repeated the analysis restricted to ileal enterocytes (n = 41,520).

### 6. Donor-stratified, pseudobulk, and cross-dataset validation

To distinguish within-cell co-variation from inter-individual (e.g., nutritional-status) differences, within-donor Spearman correlations were computed for each donor with ≥20 enterocytes; for a given gene pair, only donors in which both genes showed non-zero expression variance were retained (64–67 donors per pair), and the distribution of within-donor r was summarized by median, interquartile range, and the percentage of donors with *r* > 0. To address pseudo-replication, pseudobulk validation aggregated per-donor arithmetic mean expression; donor-level Spearman correlations were computed between the mean b^0+^ complex score (per-cell product SLC3A1 × SLC7A9, averaged per donor) and mean selenoprotein expression across the 90 of 105 donors with a non-zero mean complex score.

### 7. Software, data, and code availability

Analyses were performed in Python 3.10 using cellxgene-census (1.17.0), scanpy (1.11.5), anndata (0.11.4), pandas (2.3.3), numpy (2.2.6), scipy (1.15.3), statsmodels (0.14.6), matplotlib, and seaborn (0.13.2). The source single-cell data are publicly available through the CellxGene Census (https://cellxgene.cziscience.com); the exact dataset identifiers and query parameters are given in Supplementary Methods, making the analysis dataset fully reconstructable. All analysis and figure-generation code is openly available at https://github.com/chenlj686-beep/selenium-enterocyte-b0plus and archived at Zenodo (https://doi.org/10.5281/zenodo.20645285).

### 8. Ethical considerations

All data were obtained from publicly available, de-identified datasets in the CellxGene Census; informed consent and original data collection were performed by the originating studies. No new human-subjects research was conducted.

## Results

### 1. b^0+^ Transporter System Shows Enterocyte-Enriched Expression Peaking in the Ileum

To characterize the cell-type specificity of the b^0+^ transporter system in the human intestine, we analyzed SLC3A1 and SLC7A9 transcript-level co-expression across all major intestinal cell types. Among absorptive epithelial populations, enterocytes showed the highest b^0+^ co-expression rate (24.9%), substantially exceeding colonocytes (1.2%), BEST4+ enterocytes (1.2%), goblet cells (1.2%), Paneth cells (0.9%), tuft cells (0.5%), and transit amplifying cells (0.4%), as well as all non-epithelial cell types examined, including fibroblasts, endothelial cells, and immune cell populations (Fig. 1A). M cells of Peyer’s patches showed the highest b^0+^ co-expression rate among all intestinal cell types (35.9%); however, they were excluded from subsequent analyses because their specialized antigen-sampling transcriptional program differs fundamentally from that of absorptive enterocytes and represents a functionally distinct, non-absorptive lineage outside the scope of the present analysis of absorptive enterocyte nutrient utilization^23,24^. Large intestinal enterocytes (Enterocyte (LI)) showed markedly lower b^0+^ co-expression (0.1%) compared with small intestinal enterocytes, consistent with small-intestinal predominance with sharp small-to-large-intestinal decline (Fig. 1A).

**Figure 1.**
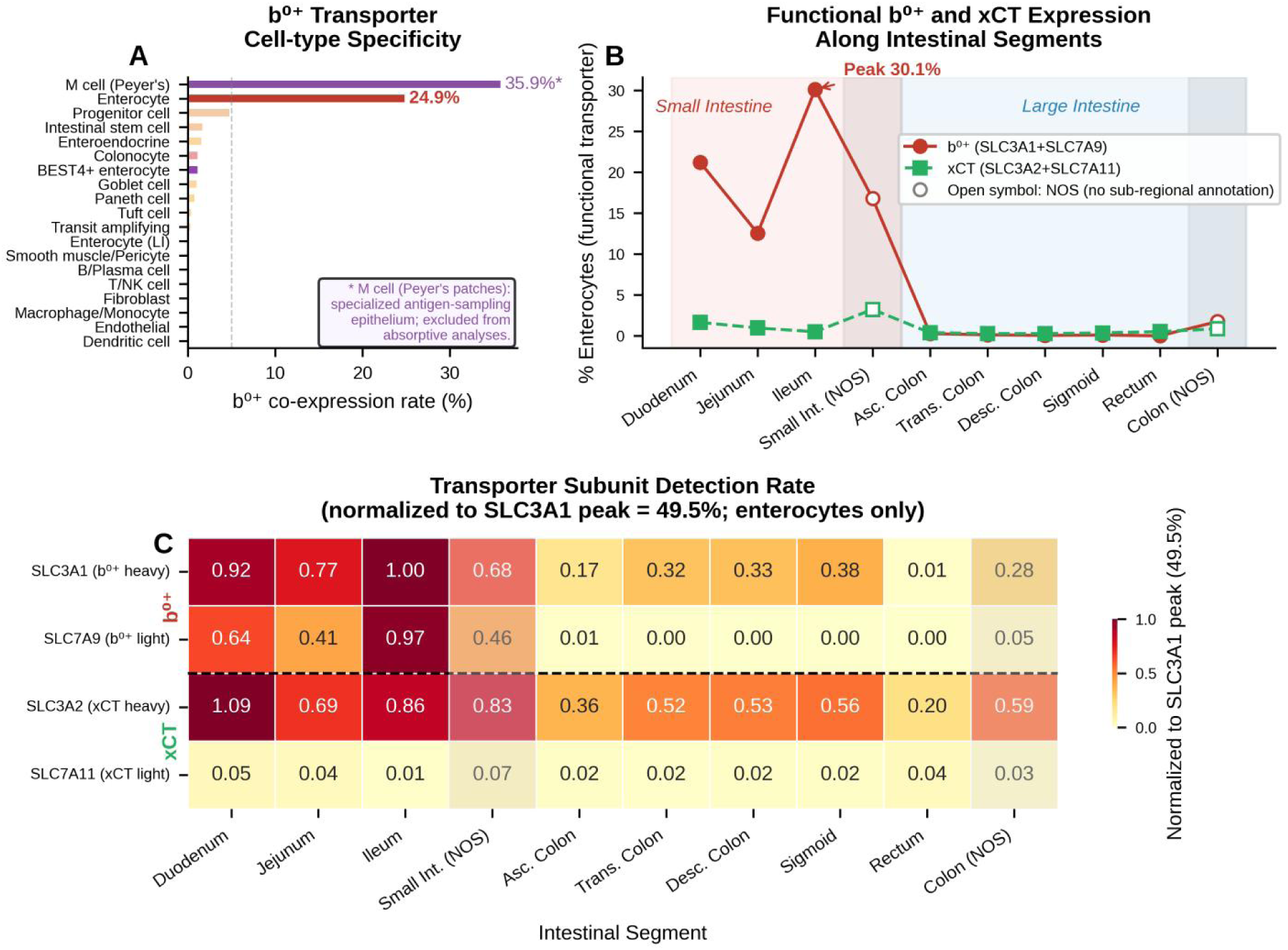
Cell-type specificity and intestinal segment distribution of the b^0+^ transporter system. (A) Bar chart showing b^0+^ (SLC3A1+SLC7A9) transcript-level co-expression rates across major intestinal cell types. Enterocytes show the highest co-expression among absorptive populations (24.9%); M cells of Peyer’s patches show the highest rate overall (35.9%) but were excluded from subsequent analyses given their non-absorptive function. (B) Line plot showing b^0+^ and xCT (SLC3A2+SLC7A11) co-expression rates along intestinal segments; b^0+^ peaks in the ileum (30.1%) and declines sharply in the large intestine. Open symbols denote segments lacking sub-regional annotation in the source dataset (NOS: not otherwise specified). (C) Heatmap showing individual detection rates of SLC3A1, SLC7A9, SLC3A2, and SLC7A11 across intestinal segments in enterocytes, normalized to SLC3A1 peak detection rate (49.5%); dashed line separates b^0+^ (upper) from xCT (lower) subunits.

Along the intestinal tract, the b^0+^ system showed a distinct spatial gradient with peak transcript-level co-expression in the ileum (30.1%), followed by the duodenum (21.2%) and jejunum (12.5%), and near-complete loss in the large intestine (mean 0.7%, p < 0.001 vs. small intestine mean 24.1%) (Fig. 1B). By contrast, the xCT system (SLC3A2/SLC7A11) maintained uniformly low transcript-level co-expression throughout all intestinal segments (mean 1.1% in small intestine, 0.5% in large intestine), remaining significantly lower than b^0+^ at every segment (p < 0.001) (Fig. 1B). At the individual subunit level, SLC3A1 and SLC7A9 detection rates showed concordant ileal enrichment and parallel decline along the large intestine (Fig. 1C). SLC3A2 maintained substantial detection rates across all segments, while its obligate partner SLC7A11 was expressed at uniformly low levels; the resulting low functional xCT co-expression likely reflects the predominant association of SLC3A2 with other light-chain partners in intestinal epithelium (Fig. 1C).

Consistent with the transcriptomic patterns identified here, publicly available protein-level data from the Human Protein Atlas indicate that SLC3A1 shows cytoplasmic and membranous expression most abundant in the small intestine, while SLC7A9 localizes to the brush border membrane of small intestinal epithelium (proteinatlas.org; reliability: Enhanced)^25^. These independent observations are compatible with the enterocyte-enriched and apically associated expression patterns observed in the present single-cell analysis.

### 2. Comprehensive Comparison of All Selenium Transporter Systems

To place the b^0+^ system in the broader context of intestinal selenium-associated transport programs, we systematically analyzed seven additional candidate transporter systems selected based on established substrate specificity or structural compatibility with distinct selenium compounds. These included transporters potentially compatible with selenomethionine (LAT1/SLC3A2/SLC7A5, LAT2/SLC3A2/SLC7A8, and ASCT2/SLC1A5), selenium oxyanion-associated transporters including DRA/SLC26A3 and PAT1/SLC26A6, selected based on the broad inorganic oxyanion exchange capacity of the SLC26 transporter family and the structural similarity of selenite to physiologic inorganic anions^26^, and candidate sulfate transporters potentially compatible with selenate (NaS1/SLC13A1 and NaS2/SLC13A4) due to the close structural similarity between sulfate and selenate^27,28^ (Fig. 2A–C).

**Figure 2.**
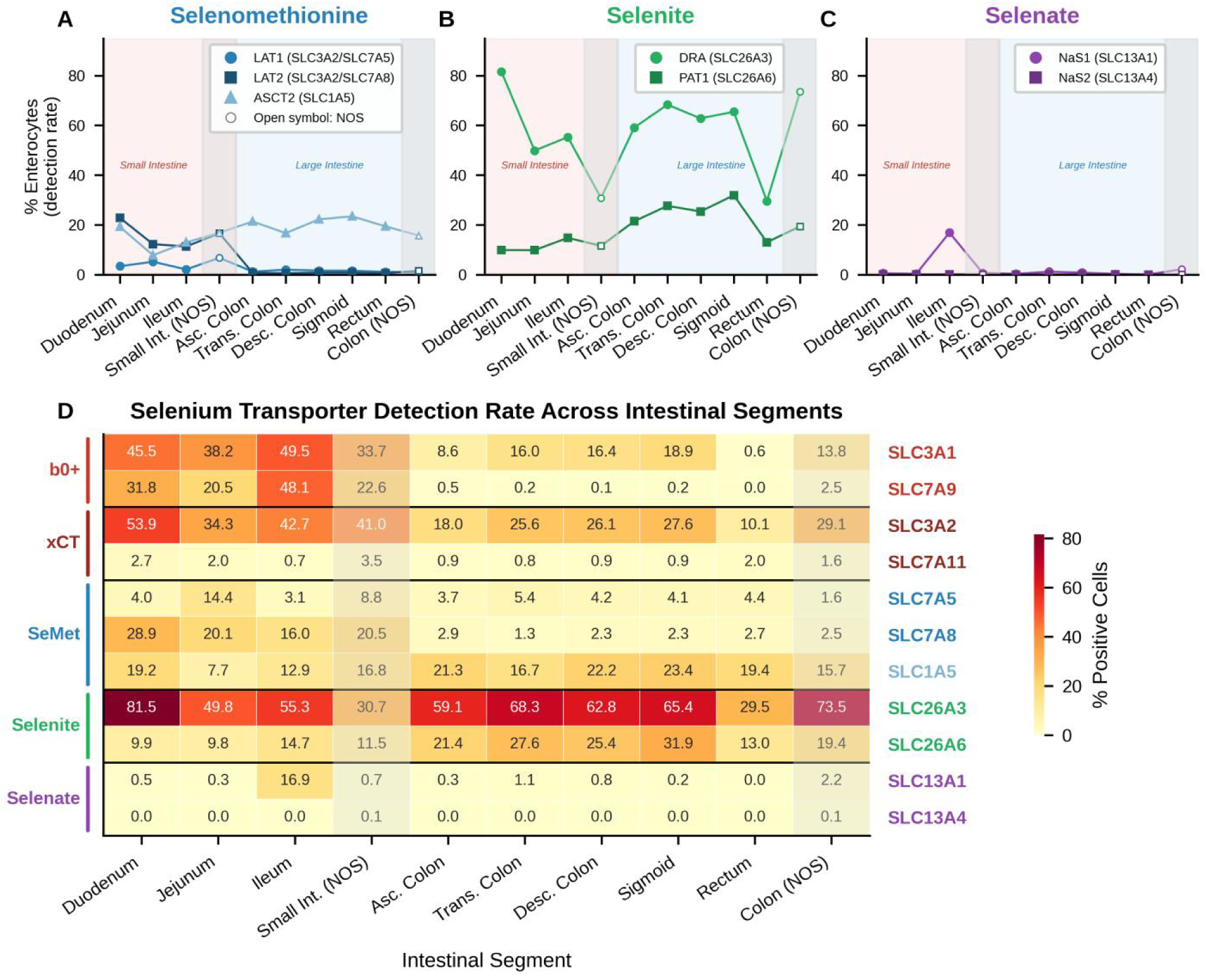
Comprehensive atlas of intestinal selenium transporter expression across all selenium forms. (A-C) Line plots showing detection rates of selenomethionine (A), selenite (B), and selenate (C) transporter systems along intestinal segments in enterocytes. Open symbols denote segments lacking sub-regional annotation (NOS). (D) Heatmap showing individual detection rates (% positive cells) for all 11 transporter subunits across intestinal segments; horizontal lines demarcate transporter system groups. Shaded columns indicate NOS segments.

Among all transporter systems examined, SLC26A3 (DRA) showed the highest overall detection rate in enterocytes across all intestinal segments (duodenum 81.5%, large intestine 59-68%), consistent with its established role as a Cl^−^/HCO₃^−^ exchanger with secondary anion transport capacity. For the selenomethionine system, SLC3A2, the common heavy chain shared by LAT1, LAT2, and xCT, showed the highest absolute detection rate among all subunits examined (duodenum 53.9%); however, its light chain partners differed markedly, with SLC7A8 (LAT2) showing substantially higher detection than SLC7A5 (LAT1) in the proximal small intestine (duodenum: 28.9% vs. 4.0%), both declining sharply in the large intestine. ASCT2 (SLC1A5) maintained relatively uniform detection across all segments (7.7-23.4%), without the proximal enrichment seen for LAT1/LAT2.

Among the selenate transporters, SLC13A4 (NaS2) was near-absent throughout the intestine (<0.1%), consistent with its primary renal expression. SLC13A1 (NaS1), however, showed an unexpected ileal-specific detection peak (16.9%), contrasting sharply with near-absent expression in the duodenum (0.5%), jejunum (0.3%), and large intestine (<3%). This ileal enrichment was independently corroborated by Human Protein Atlas immunohistochemical data, in which two independent antibodies (HPA053615 and HPA062714) both detected SLC13A1 protein in small intestinal glandular cells (Low to Medium staining intensity, cytoplasmic and membranous localization), suggesting possible ileal involvement in selenate handling, though functional validation will be required to exclude annotation artifacts ^25^.

Heatmap analysis of all 11 transporter subunits confirmed that b^0+^ exhibited the most pronounced small intestine-restricted pattern, with SLC3A1 and SLC7A9 showing concordant ileal enrichment and near-complete disappearance in the large intestine. By contrast, selenite transporters (SLC26A3/SLC26A6) maintained broad colonic expression, and SLC3A2 remained detectable throughout. These findings delineate distinct segment-specific contributions of different selenium chemical forms to intestinal selenium absorption (Fig. 2D).

Among all transporter systems examined, only b^0+^ combined small intestinal segment specificity, high functional heterodimer co-expression, and substrate preference for organic selenium forms, positioning it as the primary candidate selenium-utilization transporter program in the human intestinal epithelium.

### 3. b^0+^-Positive Enterocytes Exhibit Coordinated Selenoprotein Enrichment with a Monotonic Gradient Pattern

To examine the relationship between b^0+^ transporter expression and selenoprotein transcriptional programs, we classified enterocytes into three mutually exclusive groups based on b^0+^ complex completeness: b^0+^-complete (SLC3A1^+^/SLC7A9^+^, n = 21,234), SLC7A9-only (SLC3A1^−^/SLC7A9^+^, n = 11,874), and negative control (SLC3A1^−^/SLC7A9^−^, n = 106,961). This stratification enabled assessment of both SLC7A9 expression per se and the additional contribution of complete b^0+^ heterodimer co-expression on selenoprotein levels.

Among 24 detected selenoproteins, 21 were significantly enriched in b^0+^-complete versus negative control cells (BH-adjusted *p* < 0.001), with 18 displaying a complete monotonic gradient (b^0+^-complete > SLC7A9-only > negative control), consistent with a coordinated selenium-utilization transcriptional program associated with b^0+^ complex expression (Fig. 3A). The strongest enrichments were observed for SELENOP (detection rate 97.2% vs. 43.7% in controls; log_2_FC = 3.65), GPX4 (92.9% vs. 52.6%; log_2_FC = 2.92), MSRB1 (40.5% vs. 8.8%; log_2_FC = 3.29), SELENOS (83.6% vs. 40.7%; log_2_FC = 2.18), and SELENOK (72.4% vs. 30.2%; log_2_FC = 1.86) (Fig. 3A). In contrast, GPX1 and GPX2 did not follow this gradient. GPX1, a ubiquitously expressed cytosolic housekeeping peroxidase^29^, was broadly distributed rather than enriched within the b^0+^ program, whereas GPX2 is characteristically restricted to the crypt/proliferative compartment rather than the mature absorptive surface^30^.

**Figure 3.**
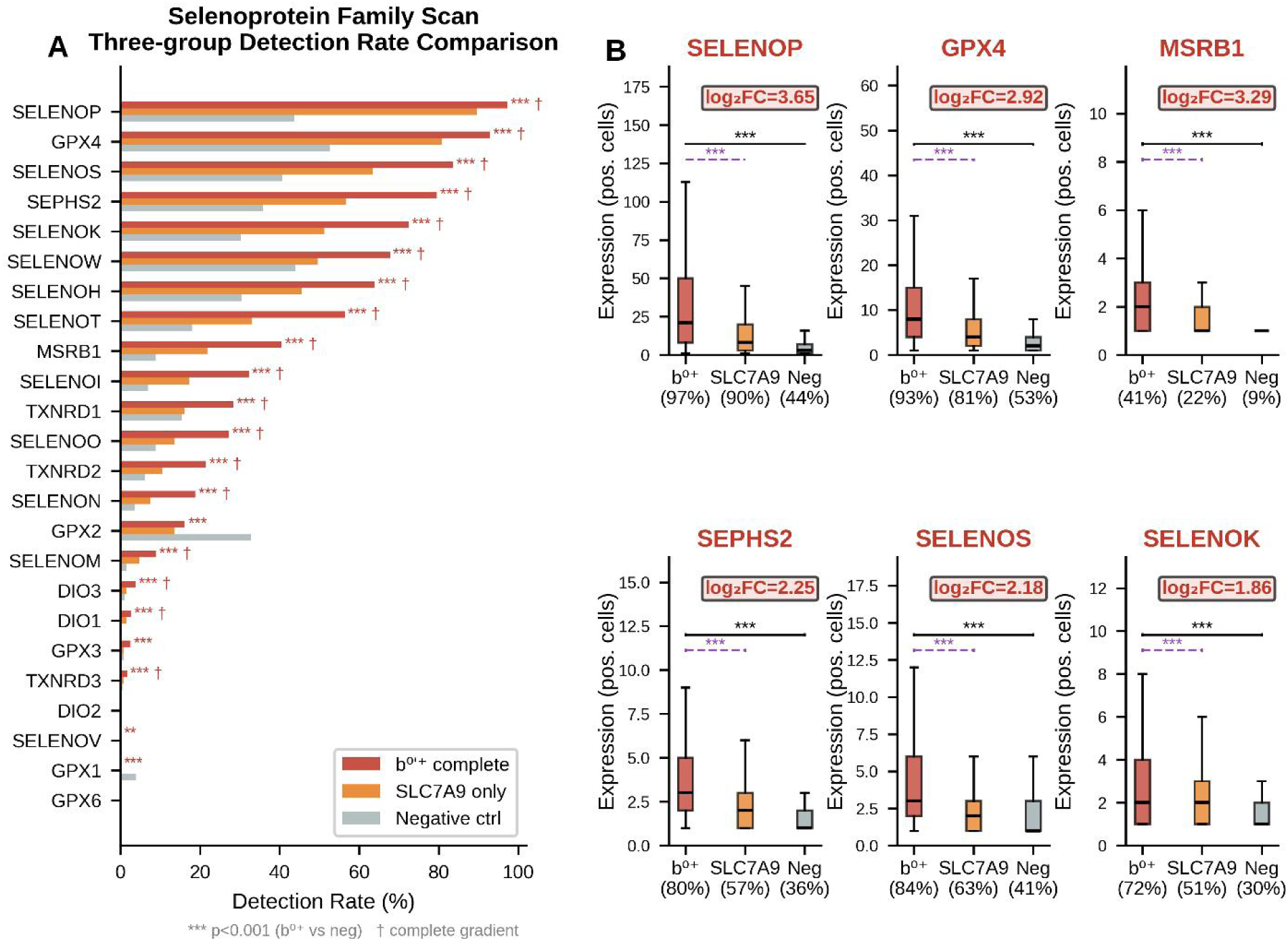
Selenoprotein enrichment in b^0+^-positive enterocytes. (A) Horizontal bar chart showing detection rates for 24 selenoproteins across b^0+^-complete (n = 21,234), SLC7A9-only (n = 11,874), and negative control (n = 106,961) enterocytes, sorted by b^0+^-complete detection rate. Asterisks indicate significance of b^0+^-complete vs. negative control comparison (*** BH-adjusted *p* < 0.001); † denotes complete monotonic gradient. (B) Boxplots showing expression distributions of six representative selenoproteins in positive cells only (expression > 0); detection rates annotated below each group. Black brackets: b^0+^-complete vs. negative control; purple dashed brackets: b^0+^-complete vs. SLC7A9-only. log_2_FC (b^0+^-complete vs. negative control) annotated in each panel.

Boxplot analysis of six representative selenoproteins confirmed that b^0+^-complete cells showed significantly higher expression than both SLC7A9-only and negative control cells (all *p* < 0.001) (Fig. 3B). Critically, b^0+^-complete cells showed additional enrichment beyond SLC7A9-only cells for all six selenoproteins (all *p* < 0.001; log_2_FC b^0+^ vs. SLC7A9-only: SELENOP 1.29, MSRB1 1.63), demonstrating that complete b^0+^ heterodimer co-expression is necessary to recapitulate the full selenoprotein transcriptional signature, and that SLC7A9 expression alone is insufficient.

### 4. The b^0+^ Transporter System Shows the Strongest and Most Maturation-independent Association with SELENOP Among All Intestinal Selenium Transporters

To assess transporter specificity, we computed Spearman correlations between selenium transporter genes and selenoproteins across 169,068 enterocytes (Fig. 4A). SLC7A9 showed the strongest associations overall (SELENOP *r* = 0.489; GPX4 *r* = 0.392; both BH-adjusted *p* < 0.001). SLC7A8 (LAT2; max *r* = 0.357) and SLC1A5 (ASCT2; max *r* = 0.349) showed intermediate correlations, while SLC7A11, SLC7A5, and SLC13A4 showed near-absent associations. SLC26A3 showed a higher-than-expected correlation with SELENOP (*r* = 0.294), likely reflecting shared mature enterocyte expression rather than selenite transport specificity.

**Figure 4.**
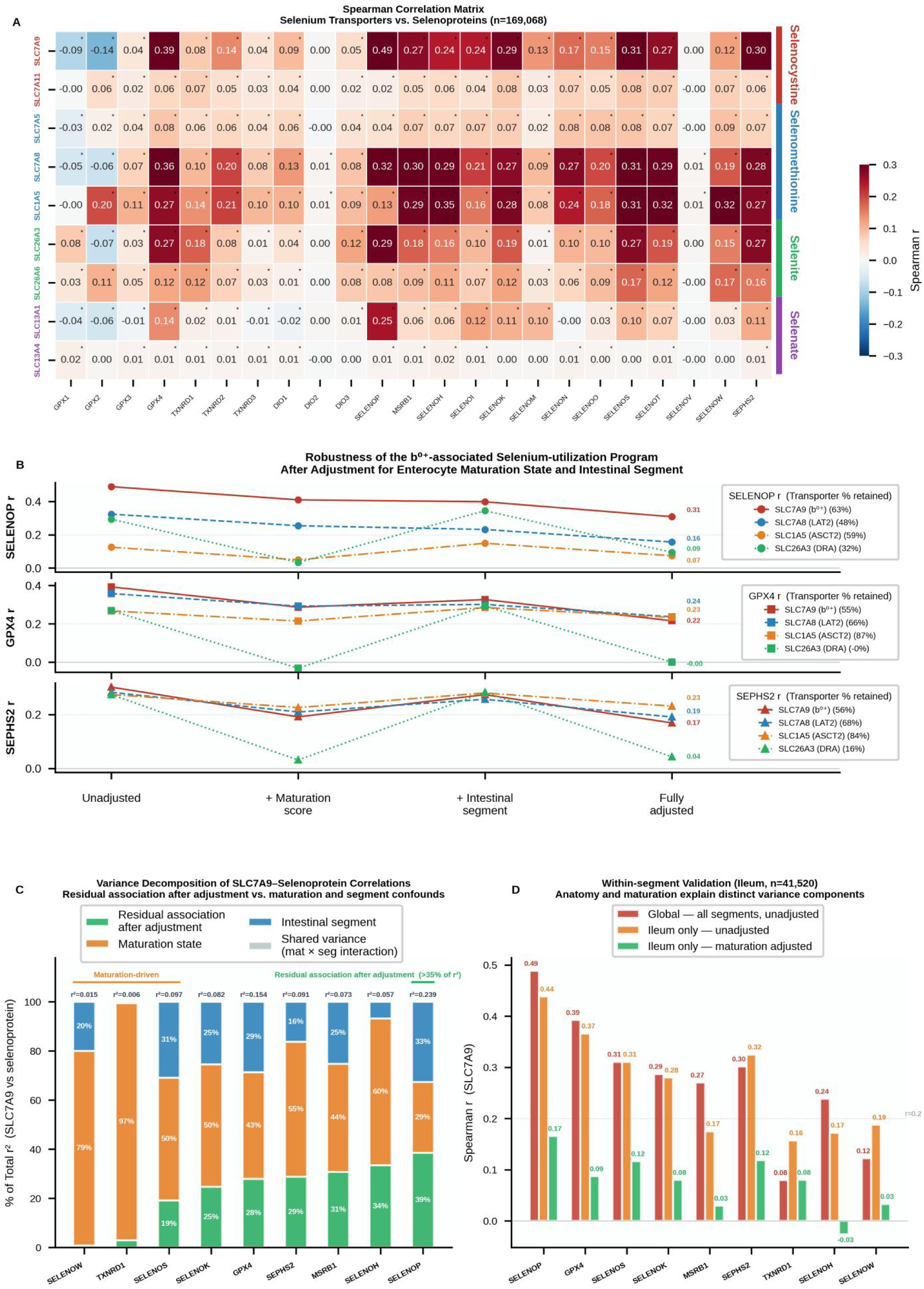
Robustness of the b^0+^-associated selenoprotein co-expression program after covariate adjustment. (A) Spearman correlation heatmap between nine selenium transporter genes and 23 selenoproteins (GPX6 excluded; constant expression) across 169,068 enterocytes; asterisks indicate BH-adjusted *p* < 0.05. Transporter labels colored by selenium form. (B) Partial Spearman correlation attenuation curves for four transporter systems (SLC7A9, SLC7A8/LAT2, SLC1A5/ASCT2, SLC26A3/DRA) against three key selenoproteins (SELENOP, GPX4, SEPHS2) under four successive covariate conditions; values indicate fully-adjusted *r* after controlling for both maturation state and intestinal segment. (C) Variance decomposition of SLC7A9-selenoprotein *r*^2^ into maturation state, intestinal segment, shared variance, and residual association components; *r*^2^ values annotated above each bar. Note: bar values represent *r*^2^ proportions; % retained values in text represent r proportions. (D) Within-ileum validation (n = 41,520) comparing global, ileum-only unadjusted, and ileum maturation-adjusted correlations.

To disentangle transporter-specific signals from enterocyte differentiation state, we computed partial Spearman correlations controlling for a 10-marker maturation score, intestinal segment, or both, across four transporters and three key selenoproteins (Fig. 4B). SLC26A3 was nearly abolished after maturation adjustment (SELENOP *r* → 0.033; GPX4 *r* → −0.032), confirming maturation co-variation as its dominant driver. SLC7A8 and SLC1A5 retained substantial independent signals for GPX4 and SEPHS2 (retention 66-87%), indicating that selenomethionine transporters contribute independent co-expression associations. However, SLC7A9 showed the highest fully-adjusted correlation with SELENOP specifically (*r* = 0.308 vs. SLC7A8 *r* = 0.156, SLC1A5 *r* = 0.074), with 39% of the total *r*^2^ retained after full adjustment versus 3% for TXNRD1 and 1% for SELENOW (Fig. 4C), and was the least attenuated of any selenoprotein after additionally controlling for sequencing depth (*r* = 0.281) (Table S1).

Within-ileum analysis (n = 41,520) confirmed that SLC7A9-selenoprotein associations were detectable within a single anatomical segment (SELENOP *r* = 0.439; GPX4 *r* = 0.366; SEPHS2 *r* = 0.325), and were partially but not fully attenuated after within-segment maturation correction (SELENOP *r* = 0.166; GPX4 *r* = 0.087; SEPHS2 *r* = 0.119), consistent with both maturation-dependent and maturation-independent components (Fig. 4D).

Taken together, while SLC7A8 and SLC1A5 also show maturation-independent selenoprotein signals, SLC7A9 (b^0+^) shows the highest fully-adjusted association with SELENOP, the primary circulating selenoprotein and biomarker of systemic selenium status, positioning b^0+^ as the dominant transporter system for the selenium-utilization program most relevant to whole-body selenium homeostasis. While the present associations are correlative, the underlying transport activity is functionally supported by our companion demonstration of SLC3A1/SLC7A9-mediated selenocystine transport, at an efficiency comparable to that of cystine, in a heterologous system^13^; the analyses below address whether this activity is coupled to a coordinated selenoprotein program at the cellular level in primary tissue.

### 5. Donor-Stratified Analysis Confirms the Specificity and Robustness of the b^0+^-Selenoprotein Association Across 105 Independent Donors

A potential confound in single-cell correlation analyses is that inter-individual variation in selenium nutritional status could generate spurious transporter-selenoprotein correlations ^31^. To address this, we performed donor-stratified Spearman correlation analysis using transporter complex co-expression scores (b^0+^: SLC7A9×SLC3A1; LAT2: SLC7A8×SLC3A2; ASCT2: SLC1A5) across donors with sufficient enterocyte representation (n ≥ 20 per donor; 105 total donors).

Within-donor analysis confirmed that b^0+^ showed the strongest and most consistent association with SELENOP (median *r* = 0.408, 100% of 64 donors *r* > 0), substantially exceeding LAT2 (median *r* = 0.172, 77.5% of 71 donors) and ASCT2 (median *r* = −0.007, 46.2% of 78 donors) (Fig. 5A). The near-zero within-donor ASCT2-SELENOP correlation, contrasting with its pooled global *r* = 0.125, corroborates the partial correlation analysis in Section 4 (both approaches agree). The b^0+^ complex also showed consistent within-donor associations across four additional selenoproteins: GPX4 (median *r* = 0.400, 100% of 65 donors), SEPHS2 (0.328, 95.4%), SELENOS (0.303, 98.4%), and SELENOK (0.273, 95.3%) (Fig. 5B).

**Figure 5.**
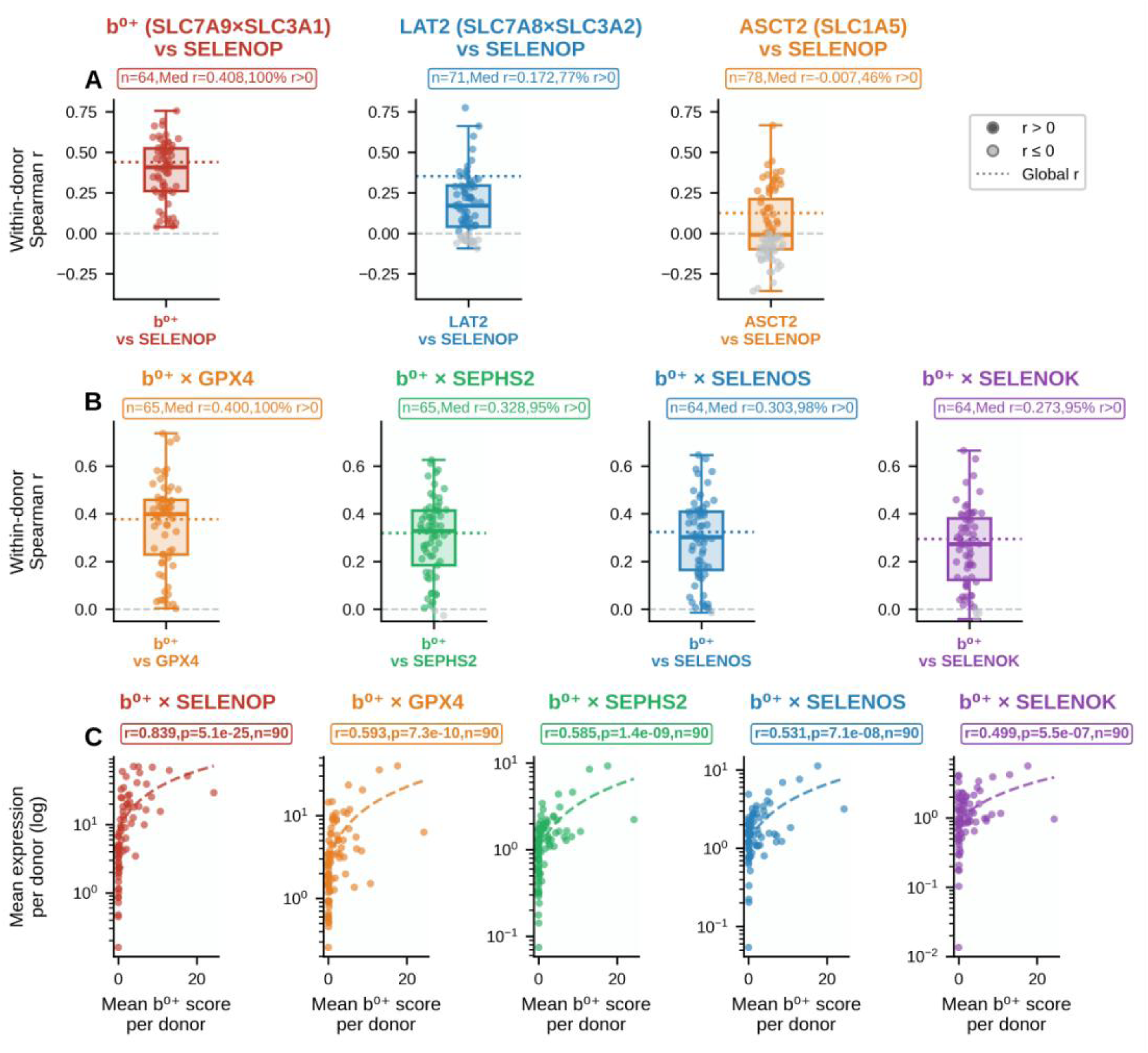
Donor-stratified and pseudobulk validation of b⁰^+^-selenoprotein associations. (A) Box-and-scatter plots comparing within-donor Spearman r for three transporter complex scores versus SELENOP; donor counts reflect those with detectable complex expression (n ≥ 20 enterocytes). Colored dots: *r* > 0; gray: *r* ≤ 0; dotted line: global *r*. (B) Within-donor b^0+^ complex correlations with four additional selenoproteins. (C) Pseudobulk validation: donor-level mean b^0+^ complex score versus mean selenoprotein expression (n = 90 donors, log scale); Spearman r and p annotated in each panel.

To further address pseudo-replication, we performed pseudobulk validation aggregating mean expression per donor (n = 90 donors with ≥ 20 enterocytes; all donors included regardless of complex score variance). Donor-level Spearman correlations between mean b^0+^ complex score and mean selenoprotein expression remained highly significant for all five pairs: SELENOP (*r* = 0.839, *p* = 5.1×10^−25^), GPX4 (*r* = 0.593, *p* = 7.3×10^−10^), SEPHS2 (*r* = 0.585, *p* = 1.4×10^−9^), SELENOS (*r* = 0.531, *p* = 7.1×10^−8^), and SELENOK (*r* = 0.499, *p* = 5.5×10^−7^) (Fig. 5C), confirming that the cell-level associations reflect genuine donor-level biological signal rather than statistical inflation from cell number.

### 6. The b0+ Enterocyte Program Co-expresses a Non-Canonical LRP Repertoire (LRP1, LRP5 and LRP6), Distinct from the Canonical SELENOP Receptors LRP2 and LRP8

To ask whether the b^0+^ program extends beyond luminal substrate import, we profiled the LRP receptors that handle SELENOP^32–35^. The intestinal repertoire was non-canonical: the prototypic extra-intestinal SELENOP receptors LRP2 (kidney) and LRP8 (brain/testis) were essentially absent from enterocytes (2.7% and 0.4% in b^0+^-complete cells), whereas LRP1, LRP5 and LRP6 were robustly expressed, markedly enriched in b^0+^-complete versus negative-control cells (50.8%, 42.5% and 24.1%; all FDR-adjusted *p* < 0.001), and present along the entire intestine (Fig. 6A, 6C).

**Figure 6.**
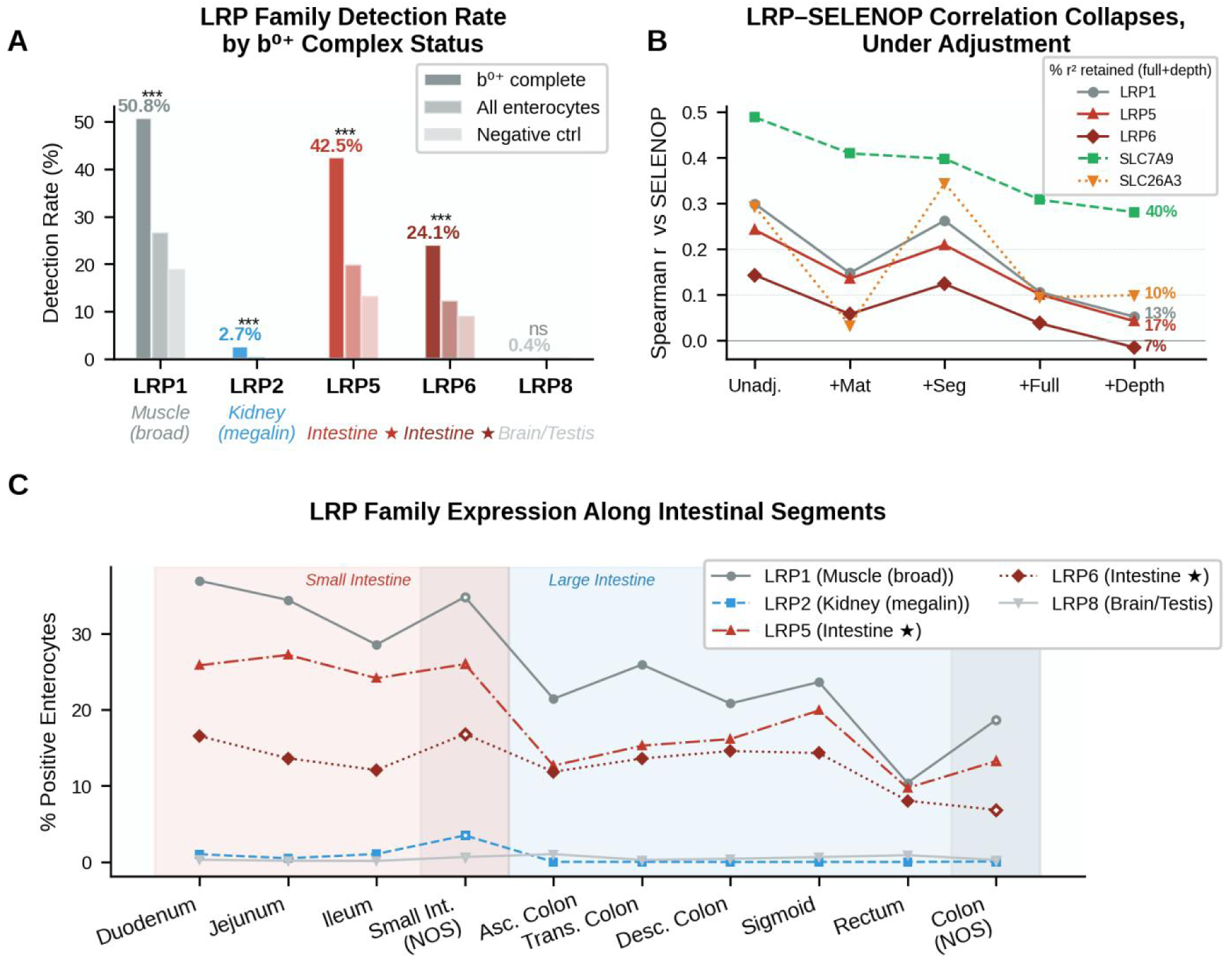
LRP family receptor expression in intestinal enterocytes. (A) Grouped bar chart showing detection rates of LRP1, LRP2, LRP5, LRP6, and LRP8 across b^0+^-complete (n = 21,234), all enterocytes (n = 169,068), and negative control (n = 106,961) groups; significance of b^0+^-complete vs. negative control comparison annotated (*** BH-adjusted *p* < 0.001; ns = not significant). Tissue-specific receptor annotations shown below x-axis. (B) Attenuation of LRP-SELENOP Spearman correlations across successive covariate adjustments (unadjusted → maturation → segment → maturation+segment → +sequencing depth). SLC7A9 (b^0+^) and the maturation-confounded exchanger SLC26A3 are shown as anchors; values at right give the percentage of raw r^2^ retained after full adjustment. (C) Detection rates of LRP family members along intestinal segments; LRP1 and LRP5 maintain broad expression throughout the intestinal tract, while LRP2 and LRP8 remain near zero at all segments.

Their correlation with SELENOP, however, was maturation-driven and unlike the b^0+^ association, did not survive adjustment. After controlling for maturation, segment and depth, LRP-SELENOP correlations fell to *r* ≤ 0.05 (7–17% of *r*^2^ retained), matching the maturation-confounded exchanger SLC26A3 and far below the 40% retained by SLC7A9 (Fig. 6B; Table S2). This contrast is itself informative: a gene set strongly co-expressed with mature enterocytes loses its SELENOP correlation under exactly the adjustment that the b^0+^ signal withstands, reinforcing that the b^0+^-selenoprotein coupling is not a generic maturation effect. A residual, above-confound co-expression with the b^0+^ complex nonetheless persisted for LRP1 and LRP5 (adjusted *r* = 0.076 and 0.108), placing these receptors within the b^0+^ enterocyte subpopulation (Table S3). Together, enterocytes carrying the b^0+^ import machinery co-express a tissue-specific, non-canonical LRP repertoire, nominating LRP1, LRP5 and LRP6 as candidate intestinal SELENOP receptors whose function awaits direct testing.

## Discussion

The present study provides a systematic single-cell characterization of nine candidate selenium transporter systems across the human intestine. Among these, the b^0+^ heterodimeric transporter (SLC3A1/SLC7A9) showed the most consistent transcriptomic association with coordinated selenoprotein enrichment across intestinal segments and donor backgrounds. Enterocytes co-expressing both subunits at the transcript level, designated b^0+^-complete cells, showed enrichment for 21 of 24 detected selenoproteins, suggesting that b^0+^ expression marks a broader selenium-utilization transcriptional state rather than a single transport activity. The apical membrane restriction of the b^0+^ system to small intestinal enterocytes and renal proximal tubule cells distinguishes it from the basolateral heavy-chain partner SLC3A2, which associates with multiple light-chain transporters including xCT^36^.

The contrasting behavior of xCT (SLC3A2/SLC7A11) in normal enterocytes is consistent with its known tissue distribution and physiological context: xCT is expressed at the basolateral membrane and is markedly upregulated in cancer to support glutathione biosynthesis and ferroptosis resistance^10,37^, whereas its expression in normal absorptive enterocytes was substantially lower and showed weaker selenoprotein association. This pattern supports the interpretation that b^0+^ and xCT represent distinct transcriptional states in intestinal epithelium, with b^0+^ marking differentiated luminal absorptive function and xCT more closely associated with stress-responsive or proliferating cell contexts^38^.

The specificity of the b^0+^-SELENOP relationship is further supported by comparison with the selenocysteine-insertion machinery. All five Sec-insertion machinery genes examined (SEPHS2, SECISBP2, EEFSEC, SEPSECS, PSTK) showed significant detection rate gradients across b^0+^-complete, SLC7A9-only, and negative control enterocytes (all p < 0.001), with SLC7A9 partial correlations retaining 53-62% of unadjusted r after full covariate adjustment (Supplementary Figure 1). However, SLC1A5 (ASCT2) and SLC7A8 (LAT2) showed substantially higher fully-adjusted correlations with these same genes (retention 83–94% and 68–81%, respectively), indicating that Sec machinery expression more broadly reflects amino acid transporter activity than a selenium-specific program. This contrasts sharply with the SELENOP result, where SLC7A9 showed the highest fully-adjusted correlation (*r* = 0.308) while SLC1A5 showed near-absent association (r = 0.074). Together, these findings suggest that b0+ expression marks a state specifically coupled to SELENOP-centered selenium utilization, whereas the broader Sec translational machinery may track a more general amino acid availability signal.

The depth-sensitivity analysis likewise argues for a genuine SELENOP-centered coupling rather than a co-detection artifact. The b^0+^-SELENOP association was the least attenuated of any selenoprotein by adjustment for sequencing depth (fully-adjusted *r* = 0.308, depth-adjusted *r* = 0.281) (Supplementary Table 1). Weaker associations eroded progressively, such as GPX4 with *r* = 0.159 and SELENOS with *r* = 0.074, and near-null pairs collapsed entirely. Such an observation runs counter to the uniform attenuation that a purely technical depth artifact would produce. Although SELENOP is predominantly synthesized by the liver, it is also expressed by intestinal epithelial cells^35,39^, where locally produced SELENOP may contribute to the enterocyte antioxidant microenvironment^6,39^. Receptor-mediated SELENOP endocytosis is tissue-specific, with LRP8, LRP2, and LRP1 mediating uptake in brain/testis, kidney, and skeletal muscle respectively^5^. In the present dataset, LRP2 and LRP8 were detected in fewer than 1% of enterocytes, while LRP1, LRP5 and LRP6 showed appreciable expression and enriched in b^0+^-complete cells. This enrichment was dissociable on adjustment: the LRP–SELENOP correlation was maturation-driven and did not survive covariate adjustment, whereas a weak but above-confound co-expression with the b^0+^ complex itself persisted (fully adjusted *r* = 0.076 and 0.108 for LRP1 and LRP5; Table S2). These receptors therefore appear to co-localize with the b^0+^ enterocyte subpopulation rather than to track SELENOP abundance per se, and we regard them as candidate receptors on the basis of this distinct, non-canonical expression pattern.Given that LRP5 and LRP6 have recently been identified as SELENOP-interacting proteins in colorectal epithelium^35^, their expression in normal primary enterocytes extends these cancer-context observations to the normal absorptive state and nominates a candidate intestinal receptor program for functional investigation.

Several important caveats apply to our findings. First, the cell-level associations reported here are transcriptomic and correlative. Although the transport function of the b^0+^ complex toward selenocystine is established in a heterologous system in our companion study^13^, the present data cannot establish that b^0+^ expression causally drives selenoprotein induction within enterocytes, as opposed to both reflecting a shared mature-absorptive transcriptional state. Demonstrating this coupling will require selenium-flux and perturbation experiments in intestinal organoid systems. Second, a substantial fraction of the SLC7A9-selenoprotein associations is attributable to enterocyte maturation state, with maturation accounting for 29-55% of explained variance for the most robustly associated selenoproteins; even after full covariate adjustment, the residual associations remain observational. Additionally, donor-stratified validation was limited to 64-67 donors with sufficient enterocyte representation (n ≥ 20 per donor) out of 105 total donors in the dataset, which may not fully capture inter-individual variability. Third, the absence of matched protein-level, spatial, or functional data limits mechanistic interpretation.

More broadly, the b^0+^ import program and this non-canonical receptor repertoire may represent two arms of a single enterocyte selenium-utilization state: at the import end, the b^0+^ exchanger supplies selenocystine substrate, demonstrated directly in our companion study^13^; at the receptor end, the same cells express an LRP repertoire for the SELENOP they produce. The far greater robustness of the b^0+^ signal to covariate adjustment is consistent with, though not proof of, b^0+^ marking the core, maturation-independent component, with the associated LRP repertoire more weakly coupled and largely maturation-scaled. Given the WNT-coreceptor role of LRP5/6 and the reported colorectal SELENOP–LRP6 interaction^35,40^, whether locally synthesized SELENOP is retained or redistributed through this receptor arm, and whether this axis links dietary selenium to WNT-governed epithelial turnover, remains an important open question that the transporter-receptor framework established here now makes experimentally tractable.

## Conclusions

This study identifies the b^0+^ transporter system (SLC3A1/SLC7A9) as the candidate selenium transporter most robustly associated with the selenoprotein-enriched transcriptional program in human intestinal epithelium. Given the well-documented prevalence of selenium deficiency in inflammatory bowel disease and colorectal cancer^41,42^, these findings provide a cellular reference framework for future mechanistic and translational studies of intestinal selenium biology.

## Data availability statement

The data presented in this study are available from the corresponding author upon reasonable request.

## Author contributions

Xiaobai He: Funding acquisition, Investigation, Methodology. Kundu Zhong and Wangzhen Yang: Investigation, Formal analysis. Jiahao Cao: Investigation. Xiaopan Chen: Writing – review & editing, Supervision. Linjie Chen: Conceptualization, Supervision, Writing –original draft and review & editing. All authors have read and approved the final manuscript.

## Acknowledgements

This work was financially supported by Zhejiang Provincial Natural Science Foundation of China under Grant No. LMS25H160004. The authors acknowledge the support from the Scientific Research Center, Hangzhou Medical College.

## Declaration on the use of generative AI tools

During the preparation of this manuscript, the authors used Claude to improve language and readability, as well as to assist with code checking and optimization. All outputs were critically reviewed and validated by the authors. The authors take full responsibility for the final content, including the correctness and integrity of the code and the overall manuscript.

## Notes

https://cellxgene.cziscience.com/census

### Competing Interest Statement

The authors have declared no competing interest.

### Summary of Updates

This revised version contains major updates to the Methods section to enhance experimental clarity and reproducibility, as well as substantial revisions to the Discussion to better contextualize the findings. Figure 4 has been reformatted for improved presentation. In addition, minor textual adjustments have been made throughout the Abstract and Results sections.

