## Supplementary information for "The b^0+^ Amino Acid Transporter Defines a Selenium-utilization Enterocyte Program in the Human Intestine"

### **Supplementary Data**

#### 1. *Supplementary Figure 1*

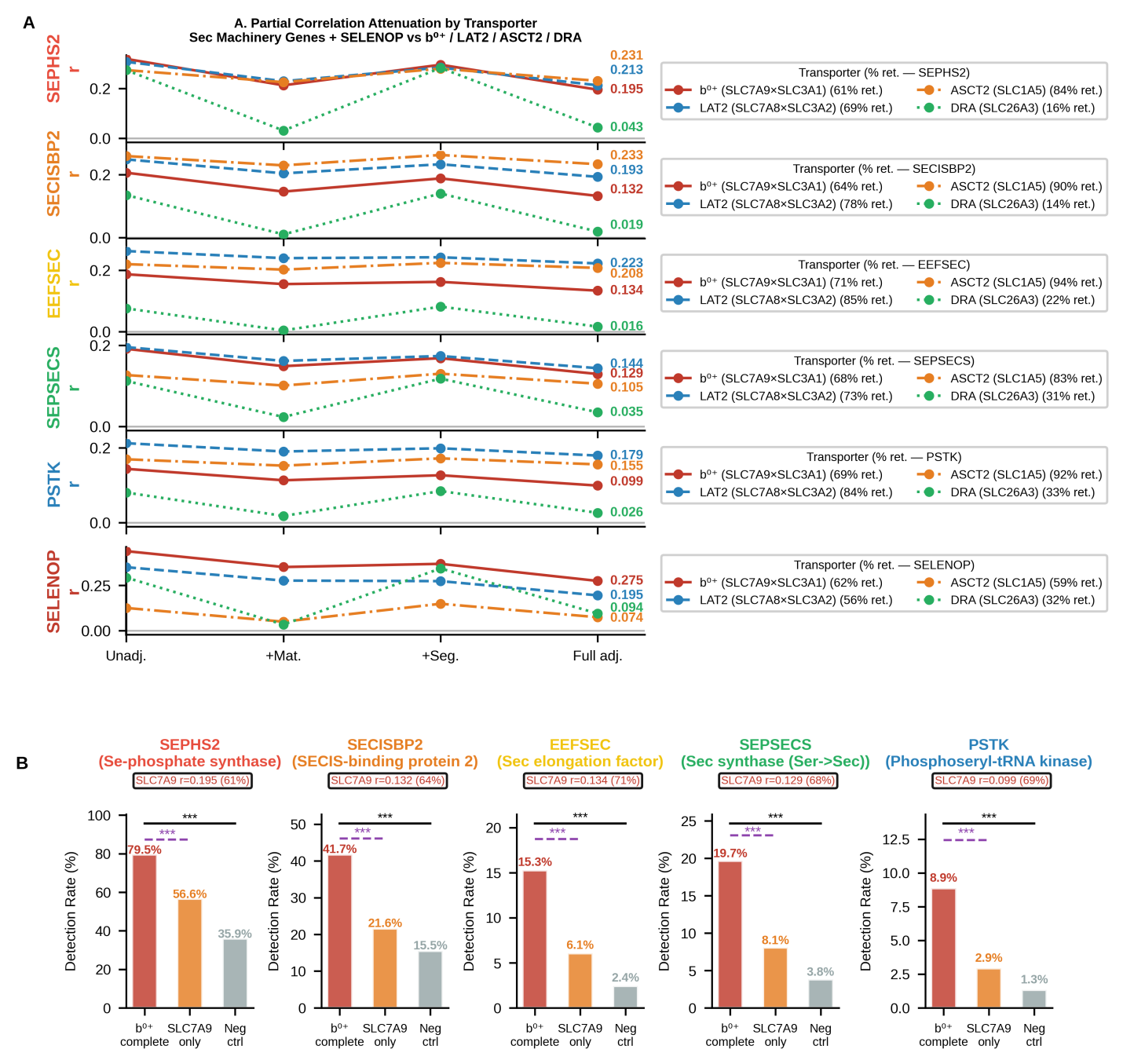

**Supplementary Figure 1. Transporter-specific partial correlation attenuation for Sec biosynthetic machinery genes and SELENOP.** (A) Partial Spearman correlation attenuation curves for four selenium transporter genes (SLC7A9/b^0+^, SLC7A8/LAT2, SLC1A5/ASCT2, SLC26A3/DRA) against five Sec-insertion machinery genes (SEPHS2, SECISBP2, EEFSEC, SEPSECS, PSTK) and SELENOP (reference, bottom row; red background) under four successive covariate conditions. Values at right indicate fully-adjusted partial r after controlling for both maturation score and intestinal segment; percentages in legend indicate retention of unadjusted r. (B) Three-group detection rates for five Sec-insertion machinery genes across b^0+^-complete (n = 21,234), SLC7A9-only (n = 11,874), and negative control (n = 106,961) enterocytes; brackets indicate pairwise significance (black: b^0+^ vs. negative control; purple dashed: b^0+^ vs. SLC7A9-only). SLC7A9 fully-adjusted partial r annotated in each panel.

### **Supplementary Methods**

#### 1. *Data Source and Cell Atlas*

Single-cell RNA sequencing data were obtained from the CellxGene Census (version 2025-11-08), a standardized, harmonized repository of publicly available human single-cell transcriptomic datasets (*cellxgene.cziscience.com*). Data were accessed programmatically via the *cellxgene-census* Python package. Three intestinal datasets were queried by dataset identifier to ensure inclusion of samples with granular intestinal segment annotation:

**Table S4** human single-cell transcriptomic datasets

| **Dataset ID** | **Description** | **Approx. Cells** |
| --- | --- | --- |
| bb0818d3-474e-44f4-a82a-169f3c62c94c ^21^ | Human Gut Cell Atlas--primary dataset | 1077238 |
| cbec7853-d996-4493-bbf8-5d82857a4e51 ^22^ | Human intestinal single-cell atlas | 263275 |
| d661aeb3-5bf7-497d-832f-2399a1ef9b63 ^21^ | Human small intestine and colon | 29498 |

Data retrieval was restricted to *Homo sapiens*, primary data only (*is_primary_data == True*). Gene expression matrices were retrieved in AnnData format (.h5ad) using *cellxgene_census.get_anndata()*, with cell metadata columns including *tissue*, *cell_type*, *dataset_id*, *assay*, and *donor_id*. All gene expression values represent raw read counts as stored in the Census.

M cells of Peyer’s patches (cell_type containing “M cell”) were excluded from all enterocyte-focused analyses a priori, as these specialized antigen-sampling epithelial cells serve distinct immunological rather than absorptive functions and are not expected to participate in dietary selenium uptake programs. This exclusion was applied prior to all downstream analyses including correlation, partial correlation, and donor-stratified validation.

#### 2. *Gene Sets*

The following gene sets were retrieved across separate query batches and merged by cell barcode:

**Selenium transporter genes (11 genes):** SLC3A1, SLC7A9 (b^0+^ system); SLC3A2, SLC7A11 (xCT); SLC7A5, SLC7A8, SLC1A5 (selenomethionine transporters); SLC26A3, SLC26A6 (selenite); SLC13A1, SLC13A4 (selenate).

**Selenoprotein family genes (24 genes):** GPX1, GPX2, GPX3, GPX4, GPX6 (glutathione peroxidases); TXNRD1, TXNRD2, TXNRD3 (thioredoxin reductases); DIO1, DIO2, DIO3 (iodothyronine deiodinases); SELENOP; MSRB1; SELENOH, SELENOI, SELENOK, SELENOM, SELENON, SELENOO, SELENOS, SELENOT, SELENOV, SELENOW; SELENOF (SEP15) was absent from the Census gene index and was therefore excluded.

**Selenocysteine insertion machinery (5 genes):** SEPHS2, SECISBP2, EEFSEC, SEPSECS, PSTK.

**Ferroptosis-related genes (2 genes):** ACSL4, LPCAT3.

**LRP receptor family (5 genes):** LRP1, LRP2, LRP5, LRP6, LRP8.

**Enterocyte maturation markers (10 genes):** ALPI, FABP1, APOA1, APOA4, SI (small intestinal absorptive markers); CA1, CA2, CEACAM5, CEACAM6, KRT20 (colonic epithelial markers). These genes were used to derive an enterocyte maturation score for partial correlation analyses (see Statistical Analysis).

All gene sets were confirmed for presence in the Census gene index prior to analysis. Gene expression was mapped from Ensembl gene IDs to HGNC symbols using the feature_name field in the var metadata.

#### 3. *Intestinal Segment Mapping*

The *tissue* field in the CellxGene Census metadata uses free-text anatomical annotations that required standardization. We mapped tissue annotations to 11 standardized intestinal segments using the following lookup table:

**Table S5** tissue annotations

| **Original tissue annotation** | **Mapped segment** |
| --- | --- |
| duodenum, duodenal epithelium | Duodenum |
| duodeno-jejunal junction | Duo-Jej Junction |
| jejunum, jejunal epithelium | Jejunum |
| ileum, ileal epithelium | Ileum |
| small intestine | Small Int. (NOS) |
| ascending colon | Ascending Colon |
| transverse colon | Transverse Colon |
| descending colon | Descending Colon |
| sigmoid colon | Sigmoid Colon |
| rectum | Rectum |
| colon, colonic epithelium | Colon (NOS) |

Cells without a mappable tissue annotation were excluded from segment-level analyses. The small intestine was defined as Duodenum, Duo-Jej Junction, Jejunum, Ileum, and Small Int. (NOS); the large intestine as Ascending Colon through Colon (NOS). The “Small Int. (NOS)” and “Colon (NOS)” categories represent cells whose tissue of origin was recorded without segment-level resolution and are treated separately in segment gradient analyses.

#### 4. *Transporter System Classification*

Selenium transporter systems were classified according to their known biochemical subunit composition and selenium substrate specificity as follows:

**Table S6** Selenium transporter systems

| **System** | **Heavy chain** | **Light chain** | **Type** | **Reported Selenium substrate** |
| --- | --- | --- | --- | --- |
| b^0+^ | SLC3A1 | SLC7A9 | Heterodimer | Selenocystine |
| xCT | SLC3A2 | SLC7A11 | Heterodimer | Selenocystine |
| LAT1 | SLC3A2 | SLC7A5 | Heterodimer | Selenomethionine |
| LAT2 | SLC3A2 | SLC7A8 | Heterodimer | Selenomethionine |
| ASCT2 | - | SLC1A5 | Monomer | Selenomethionine |
| DRA | - | SLC26A3 | Monomer | Selenite |
| PAT1 | - | SLC26A6 | Monomer | Selenite |
| NaS1 | - | SLC13A1 | Monomer | Selenate |
| NaS2 | - | SLC13A4 | Monomer | Selenate |

For heterodimeric systems, transcript-level co-expression was defined as simultaneous detection of both the heavy chain and light chain subunit (expression count > 0). For monomeric transporters, detection of the single subunit gene was used directly. The SLC3A2 heavy chain is shared among xCT, LAT1, and LAT2 systems; cells were independently classified for each system based on the respective light chain.

*5. Single-dataset effects analysis.*

To exclude single-dataset effects, Spearman correlations between the b^0+^ complex score (SLC3A1 × SLC7A9) and each selenoprotein were recomputed independently within each contributing dataset (n = 117,878; 44,893; 6,297 enterocytes). Direction was consistent across all three datasets for 15 of 24 selenoproteins; five genes (DIO1, DIO2, DIO3, SELENOV, GPX6) had near-zero or undefined r values precluding direction assessment; SELENOH and SELENOI showed negative *r* in the Xu et al. dataset, likely reflecting its colonic tissue enrichment (see Table S7 footnote). Full results are provided in Table S7.

*6. Sequencing-depth sensitivity analysis.*

Because both gene co-detection and selenoprotein detection scale with per-cell capture efficiency, sequencing depth was tested as a potential confound. The log-transformed number of non-zero detected genes per cell, log(nnz + 1), from the Census cell metadata, was added as a third covariate to the partial-correlation model, and to the three-group comparisons (by regressing log(nnz + 1) out of expression ranks before computing rank-biserial effect sizes). Depth-adjusted partial correlations were compared with their fully-adjusted counterparts to quantify the depth-attributable fraction of each association.

#### *7. Group Classification and Co-expression Statistics.*

#### Complete co-expression statistics for the SLC3A1/SLC7A9 system, including per-group cell counts and detection rates, conditional probabilities, Spearman correlation, threshold sensitivity analysis (count > 0, ≥2, ≥3), and pairwise sequencing-depth comparisons across the three analysis groups, are provided in Supplementary Table S8.

#### *8. Visualization*

All visualizations were generated in Python using *matplotlib* and *seaborn*. Gene expression heatmaps used row-normalization (each row divided by its maximum value) to enable cross-segment pattern comparison while preserving within-row directionality. Absolute detection rate heatmaps used the YlOrRd colormap; normalized heatmaps used Blues. Spearman correlation heatmaps used the RdBu_r diverging colormap centered at zero (range −0.3 to +0.3). All figures were saved at 300 dpi minimum.
